# Silver nanoparticles induce a *triclosan-like* antibacterial action mechanism in multi-drug resistant *Klebsiella pneumoniae*

**DOI:** 10.1101/2020.10.24.337204

**Authors:** Vikram Pareek, Stéphanie Devineau, Sathesh K. Sivasankaran, Arpit Bhargava, Jitendra Panwar, Shabarinath Srikumar, Séamus Fanning

**Affiliations:** UCD-Centre for Food Safety, UCD School of Public Health, Physiotherapy and Sports Science, University College Dublin, Dublin D04 V1W8, Ireland; Department of Biological Sciences, Birla Institute of Technology & Science, Pilani 333031, India; Université de Paris, BFA, UMR 8251, CNRS, F-75013 Paris, France; Genome Informatics Facility, Iowa State University, 448 Bessy Hall, Ames, IA, 50011, USA; Department of Food, Nutrition and Health, College of Food and Agriculture, UAE University, Al Ain, UAE; Institute for Global Food Security, Queen’s University Belfast, 19 Chlorine Gardens, Belfast BT9 5DL, United Kingdom

**Keywords:** *Klebsiella pneumoniae*, silver nanoparticles, RNA sequencing, soxS, triclosan

## Abstract

Infections associated with antimicrobial-resistant bacteria now represent a significant threat to human health using conventional therapy, necessitating the development of alternate and more effective antibacterial compounds. Silver nanoparticles (Ag NPs) have been proposed as potential antimicrobial agents to combat infections. A complete understanding of their antimicrobial activity is required before these molecules can be used in therapy. Lysozyme coated Ag NPs were synthesized and characterized by TEM-EDS, XRD, UV-vis, FTIR spectroscopy, zeta potential and oxidative potential assay. Biochemical assays and deep level transcriptional analysis using RNA sequencing were used to decipher how Ag NPs exert their antibacterial action against multi-drug resistant *Klebsiella pneumoniae* MGH78578. RNAseq data revealed that Ag NPs induced a *triclosan-like* bactericidal mechanism responsible for the inhibition of the type II fatty acid biosynthesis. Additionally, released Ag^+^ generated oxidative stress both extra- and intracellularly in *K. pneumoniae*. The data showed that *triclosan-like* activity and oxidative stress cumulatively underpinned the antibacterial activity of Ag NPs. This result was confirmed by the analysis of the bactericidal effect of Ag NPs against the isogenic *K. pneumoniae* MGH78578 *ΔsoxS* mutant, which exhibits a compromised oxidative stress response compared to the wild type. Silver nanoparticles induce a *triclosan-like* antibacterial action mechanism in multi-drug resistant *K. pneumoniae*. This study extends our understanding of anti-*Klebsiella* mechanisms associated with exposure to Ag NPs. This allowed us to model how bacteria might develop resistance against silver nanoparticles, should the latter be used in therapy.

## Introduction

Antimicrobial resistance (AMR) is responsible for approximately 700,000 deaths annually across the globe and this number is expected to increase further if new measures are not adopted and antibacterial compounds discovered.(1) The emergence of multi-drug resistance (MDR) in various pathogenic bacterial species represents a serious public health challenge,(2, 3) leading to hospital- and community-acquired infections, which are difficult to treat and control.(4, 5) Since AMR is estimated to overtake cancer as the main cause of death in 50 years, innovative approaches including the development of novel antimicrobial strategies using silver nanoparticles (Ag NPs) are required.(6, 7)

*Klebsiella pneumoniae* is one of the members of the ESKAPE pathogens (representing *Enterococcus faecium, Staphylococcus aureus, Klebsiella pneumoniae, Acinetobacter baumannii, Pseudomonas aeruginosa* and *Enterobacter* spp.). (4) It is a member of Enterobacteriaceae family - Gram-negative, non-motile, and rod-shaped. This bacterium is considered an opportunistic pathogen commonly found in the intestine, mouth, and skin of humans. It is mainly associated with hospital-acquired infections (nosocomial infections) and responsible for respiratory/urinary tract infections, pneumonia, and sepsis.(1, 8, 9) This pathogen can also form biofilms on indwelling medical devices leading to persistent nosocomial infection.(10, 11) About 25 % of nosocomial *K. pneumoniae* were found to be resistant to carbapenem-based compounds.(12, 13) *K. pneumoniae* were also found to be resistant to colistin, a last-line antibiotic.(14, 15) The emergence of AMR in *K. pneumoniae* against critically important classes of antibiotics represents a major threat to conventional clinical therapy.(11, 16) Novel antibacterial strategies are required to overcome this challenge.(17, 18)

Silver and other metals such as copper and zinc have historically been used as potential antibacterial agents.(19, 20) Antibacterial activity of silver can vary depending on its chemical form.(21–23) Metallic forms continuously release small numbers of ions, which makes it a slow-acting agent, whilst the ionic form is more efficient. Although Ag^+^ is reported to exhibit better antibacterial activity,(24) direct exposure to mammalian cells has toxic side effects that limit its application in therapy.(25, 26) In contrast, Ag NPs provide a greater surface area, leading to a more controlled release of Ag^+^.(27) Hence, this form has potential as an antibacterial compound.(28, 29) Despite this advantage, little is known about the antibacterial action mechanisms of Ag NP based formulations.

We synthesized lysozyme coated Ag NPs (L-Ag NPs) and characterized their antibacterial mechanism against MDR *K. pneumonia* MGH78578 using chemical analysis, biochemical assays and deep-level RNA sequencing. Our data revealed that L-Ag NPs induced a *triclosan-like* antibacterial effect against MDR *K. pneumoniae.* The inhibition of the type II fatty acid biosynthesis along with Ag^+^ induced oxidative stress were responsible for the anti-*K. pneumoniae* effect. Our results provide molecular insights into how bacteria might deploy antibacterial strategies to counteract the toxic effects of Ag NPs. This allowed us to model how *K. pneumoniae* might develop resistance against Ag NPs.(30)

## Results and discussion

### Synthesis and characterization of L-Ag NPs

L-Ag NPs were synthesized by a co-reduction method wherein lysozyme functions both as a reducing and stabilizing agent using heat reflux action at 120 °C.(31, 32) Lysozyme acts as a capping material devoid of enzymatic activity. We selected L-Ag NPs based on their bactericidal activity against MDR *K. pneumoniae* MGH78578. The synthesis of L-Ag NPs was associated with the appearance of the plasmon peak at 414 nm (Figure 1A).(33) The average diameter of L-Ag NPs measured by TEM was 5.2 ± 1.2 nm (Figures 1B-C). The X-ray diffraction pattern obtained corresponds to the face-centered cubic lattice structure of crystalline silver (JCPDS file 04-0783) (Figure 1D, S1).(34, 35) The surface charge of L-Ag NPs was estimated by measuring their zeta potential in water and modified LB (mLB) medium (devoid of NaCl) at 37° C. L-Ag NPs were found to be negatively charged in both conditions with ζ = −38.2 ± 1.6 mV in water and ζ = −24.2 ± 0.9 mV in mLB medium. The 1,632 cm^−1^ peak in the FTIR spectrum corresponds to the amide I vibration characteristic of the protein backbone, confirming the presence of lysozyme associated with Ag NPs (Figure 1E).(36)

**Figure 1.**
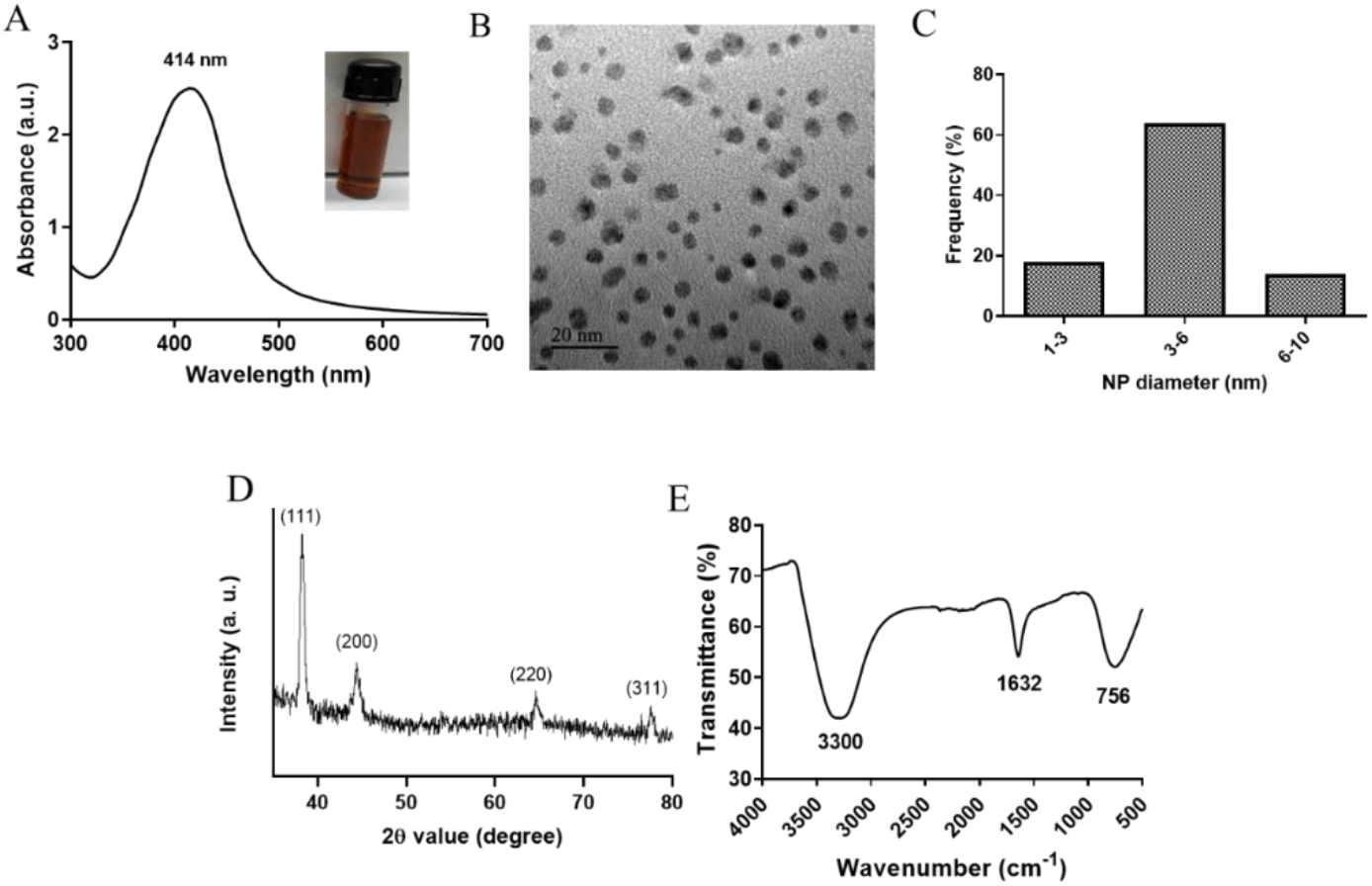
Characterization of L-Ag NPs. (A) UV-visible spectrum (inset shows a picture of the L-Ag NP suspension after synthesis). (B) TEM image. (C) Distribution of L-Ag NP diameter determined from TEM analysis. (D) XRD diffraction spectrum. (E) FTIR spectrum.

### L-Ag NPs inhibit the proliferation of *K. pneumoniae* MGH78578

In media with high salt content like chloride and phosphate, Ag NPs may aggregate and free Ag^+^ can precipitate, potentially reducing their bactericidal activity.(37–40) Hence, the bactericidal effect of L-Ag NPs against *K. pneumoniae* MGH78578 was assessed in mLB medium (LB medium devoid of NaCl).(41, 42) No significant difference between the growth of *K. pneumoniae* in mLB and LB media was observed (Figure 2A) confirming mLB had no phenotypic effect. Approximately 3×10^7^ CFU mL^−1^ log_10_ phase bacterial cells were exposed to L-Ag NPs. The MIC of L-Ag NPs was 21 μg (Ag) mL^−1^ (Figure 2B). To determine the bactericidal efficiency of L-Ag NPs, *K. pneumoniae* MGH78578 exposed to L-Ag NPs were spread plated and incubated for 12 h at 37 °C. The MBC of L-Ag NPs was 45 μg (Ag) mL^−1^ (Figure S2). These results show that L-Ag NPs inhibit the proliferation of *K. pneumoniae* MGH78578 at low concentrations.

**Figure 2.**
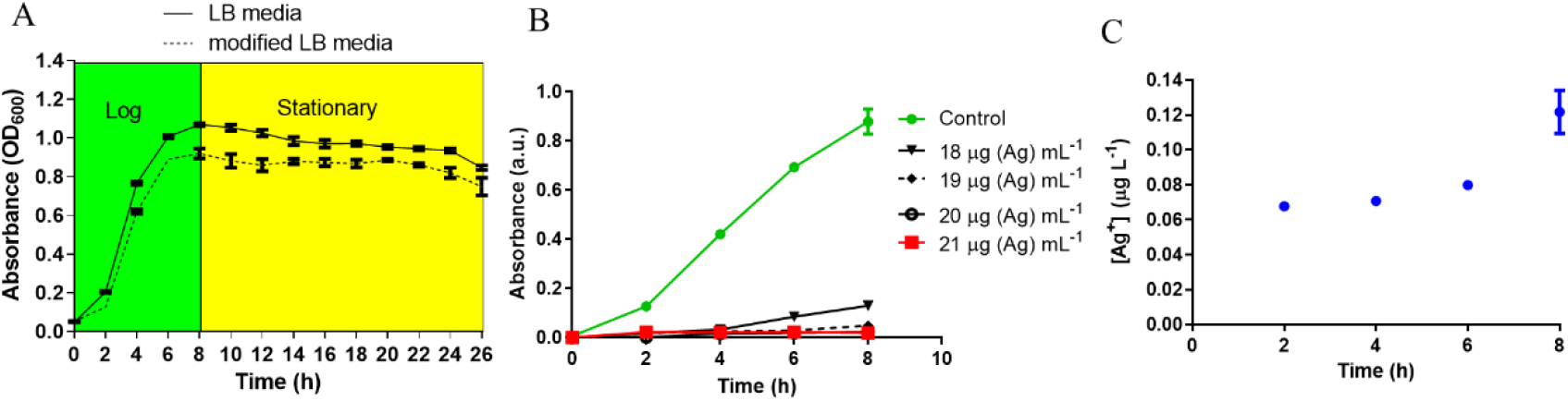
(A) Bacterial growth curve analysis of *K. pneumoniae* MGH78578 in LB and mLB media. (B) Determination of *K. pneumoniae* L-Ag NP MIC in mLB medium. (C) Dissolution kinetics of L-Ag NPs in mLB medium at 37°C.

### L-Ag NPs generate ROS and limited membrane damage in *K. pneumoniae* MGH78578

Reactive oxygen species (ROS) could be generated outside the bacterial cells by L-Ag NPs and released Ag^+^.(43) Limited dissolution of L-Ag NPs was observed in mLB media for 8 h at 37°C (Figure 2C). Extracellular ROS production was evaluated by measuring the oxidative potential of L-Ag NPs in a simplified synthetic respiratory tract lining fluid. The depletion of antioxidants (uric acid, acetic acid, and reduced glutathione GSH) was also noted when measured by HPLC after 4 h incubation at 37 °C (Figure 3A).(44) We observed a dose-dependent depletion in ascorbic acid and GSH (up to 100%), showing that, at low NP concentration, L-Ag NPs can generate ROS extracellularly.

**Figure 3.**
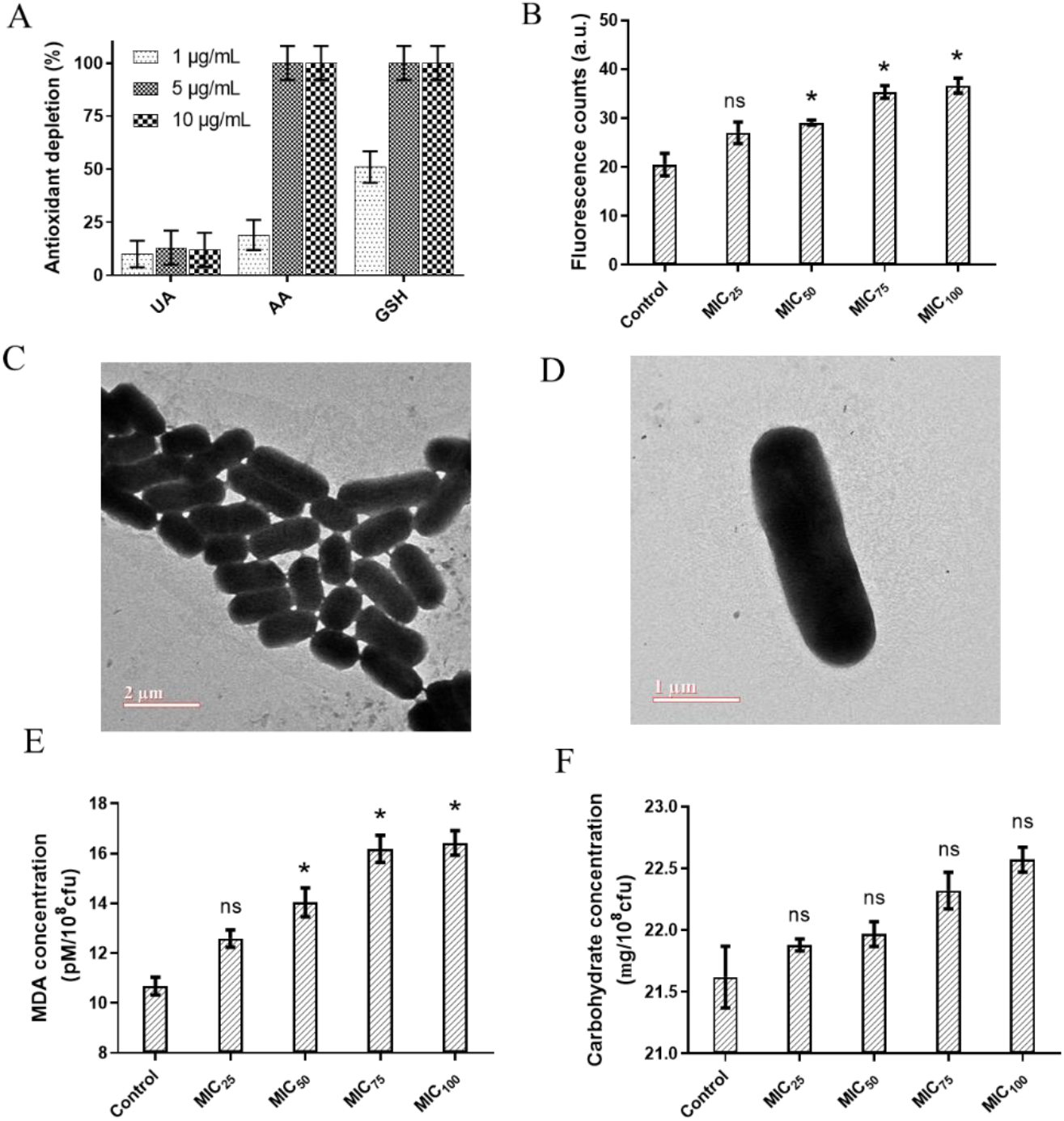
(A) Acellular oxidative potential of L-Ag NPs measured by the depletion of antioxidants, uric acid (UA), ascorbic acid (AA), and reduced glutathione (GSH), in simplified synthetic lung lining fluid at 37°C. (B) Analysis of the intracellular ROS generation in *K. pneumoniae* MGH78578 following exposure to L-Ag NPs by the DCFH-DA assay. (C,D) Representative TEM images of untreated *K. pneumoniae* (control) (C) and *K. pneumoniae* exposed to MIC75 of L-Ag NPs (D). (E,F) Membrane damage analysis of *K. pneumoniae* following exposure to L-Ag NPs. Analysis by MDA assay (E) and anthrone assay (F).

Then, we measured the concentration of silver that enter *K. pneumoniae* MGH78578 by ICP-OES following treatment with L-Ag NPs for 5, 30, and 60 min. The uptake of silver, either in the form of L-Ag NPs or Ag^+^, was proportional to the time duration of treatment (Figure S3). To determine whether ROS were also produced inside the bacterial cells following exposure to L-Ag NPs, we used the DCFH-DA assay.(45) A dose-dependent increase in the intracellular ROS was observed, a reflection of the oxidative stress external to the bacterial cell (Figure 3B).

The severe oxidative stress induced by L-Ag NPs could lead to many downstream phenotypes in bacteria including compromised membrane integrity.(23, 27, 28, 46) TEM analysis of *K. pneumoniae* did not show any change in the membrane integrity of the exposed bacterial cells (Figures 3C-D). To verify this observation, we explored the degree of bacterial membrane damage in the whole population using an MDA assay. A limited though significant concentration-dependent increase in the level of lipid peroxidation was observed (Figure 3E). Additionally, a concentration-dependent increase of carbohydrate release was observed following L-Ag NP exposure (Figure 3F). Overall, our results show that L-Ag NPs induced limited bacterial membrane damage, a phenotype that was not observed by TEM.

### Transcriptomic profiling of L-Ag NPs exposed *K. pneumoniae* MGH78578 using RNA-seq

We used RNA-seq to understand how *K. pneumoniae* MGH78578 responded to L-Ag NP exposure following a previously standardized protocol investigating the transcriptome of *K. pneumoniae* MGH78578 exposed to sub-inhibitory concentrations of a chemosensitizer.(47) Here, we exposed *K. pneumoniae* MGH78578 to a sub-inhibitory concentration of L-Ag NPs at MIC_75_ (15.8 μg (Ag) mL^−1^) for a period of 5- and 30-min. The 5 min time point was used to identify the transcriptional signals associated with early exposure, whilst the 30 min time point demonstrated the adaptive responses of L-Ag NP exposed *K. pneumoniae* (replication time is 20 min). Approximately 336 million reads were obtained across all 8 libraries with an average of 42 million reads *per* library (Table S1) sufficient for downstream transcriptomic analysis.(47, 48) The *VOOM* function in the *limma* package(49) was used to identify differentially regulated genes. We obtained statistically significant data (p ≤ 0.05) for a total of 3,760 and 3,945 genes following 5 min and 30 min exposure respectively (Figure 4A) (Dataset WS1). Statistically significant genes with a log2 fold change expression of ≥ 1.0 and ≤ −1.0 (exposed *versus* unexposed cells) were considered to be up- and down-regulated respectively.(47, 50) For the post-5 min exposure, 1,008 genes were found to be upregulated and 1,063 genes were downregulated. At 30 min post exposure, 1,205 genes were found to be upregulated and 1,246 genes were downregulated (Figure 4B). Among all the upregulated genes at the 5- and 30-min, 705 genes were found to be commonly upregulated, whereas 303 and 500 genes were uniquely upregulated at 5 min and 30 min respectively. For downregulated genes, 810 genes were found to be commonly downregulated at both time points, whereas 253 and 436 genes were specifically downregulated at 5 min and 30 min (Figure 4C).

**Figure 4.**
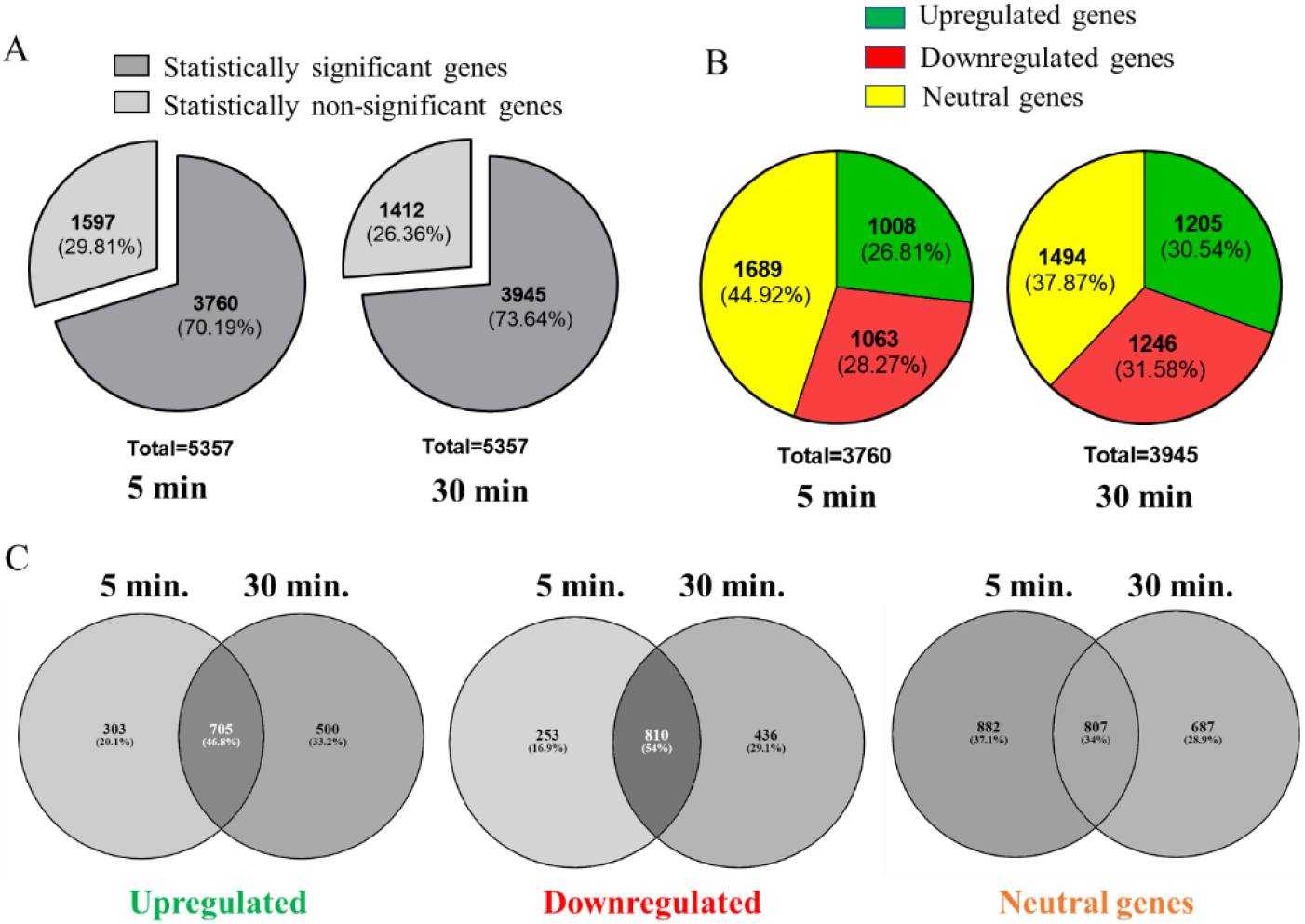
Overview of the RNAseq analysis of *K. pneumoniae* MGH78578 following treatment with MIC_75_ L-Ag NPs. (A) Statistically significant and insignificant genes recorded at 5 and 30 min post-treatment. (B) Statistically significant upregulated, downregulated and neutral genes identified at 5 and 30 min post-treatment. (C) Comparison of the upregulated, downregulated, and neutral genes identified at 5 and 30 min post-treatment.

We selected representative genes related to the bacterial defence system (*soxS*), transcriptional regulator (*ramA*), outer membrane porin proteins (*ompC*) and virulence (*rfaH*) to validate our transcriptomic data using qRT-PCR. Gene expression analysis reflected a similar pattern of results when compared to RNA-seq data (Figure S4).

### Bacterial oxidative stress response following exposure to L-Ag NP

Our RNA-seq data shows that the oxidative stress response induced by ROS generation is choreographed by both *soxS* and *oxyR,* two major transcriptional factors that respond to oxidative stress in Enterobacteriaceae.(51) The OxyR system functions as a global regulator of peroxide stress whilst the SoxRS system is involved in the control of superoxide stress.(51, 52) Increased intracellular ROS levels induce the oxidation of OxyR protein, which in turn activates detoxifying processes including heme biosynthesis, thiol-disulfide isomerization, among others.(53) In the case of the SoxRS regulon, oxidative stress causes the oxidization of the SoxR protein, which activates the *soxS* transcriptional regulator, thereby triggering a bacterial defense mechanism involving various efflux pumps and redox counter measures.(51) In L-Ag NP exposed *K. pneumoniae* MGH78578, *soxS* was highly upregulated (300- and 88-fold in 5 and 30 min) and in contrast *oxyR* was moderately upregulated (approximately 5-fold in both time points). This suggested that the oxidative stress regulon was active throughout the 30 min of L-Ag NP exposure time, particularly for the *soxS* regulon.

To confirm this, we compared our RNA-seq dataset to the *K. pneumoniae* MGH78578 <u>oxidative *soxS* regulon</u> generated earlier.(54) We found that of the 254 genes belonging to the <u>oxidative *soxS* regulon</u>, 129 (51%) had similar expression patterns in both datasets, confirming that the *soxS* regulon is activated in L-Ag NP exposed *K. pneumoniae* (Dataset WS2). A typical example of *soxS* induction can be visualized in the expression of *fpr* [ferredoxin-NADP^+^ reductase], which was induced at 22- and 9-fold at 5 and 30 min. In response to *soxS* activation, a redox neutralization process is triggered related to the overexpression of *fpr* by 4.5-fold at 5 min, that then reduced to 3.2-fold at 30 min post-treatment.(55)

### Activation of efflux pumps following exposure to L-Ag NP

Bacteria activate various efflux pumps to expel the toxic Ag^+^.(56, 57) Classic efflux pump-encoding genes such as *acrAB-tolC* (58, 59) and *marRAB* were found to be highly upregulated following treatment with L-Ag NPs, a feature that was sustained throughout the testing period of 30 min, possibly driven by the positive regulation *via* SoxS.(51, 54) Another efflux pump encoded by the *silCFBA* operon conferring a silver resistance phenotype was also upregulated.(57, 60) Transcription of the *silCFBA* operon and *silP* are controlled by the *silRS*, which encodes a two-component system wherein SilR acts as a response regulator and SilS acts as a histidine kinase.(61, 62) SilP is a P-type ATPase efflux pump, which facilitates the passage of Ag^+^ from cytoplasm to the periplasm. SilF acts as a chaperone and transfers Ag^+^ from periplasm to the SilCBA complex, a three-protein dependent cation/proton antiporter system. Another protein from silver resistance system is SilE, present downstream to the *silRS* (Figure 5).(61, 62) Upregulation of *silC/B/E/R/S/P* and KPN_pKPN3p05946 genes were noted in our RNA-seq data showing the activation of efflux pumps, encoding a possible silver resistance in response to L-Ag NP exposure.

**Figure 5.**
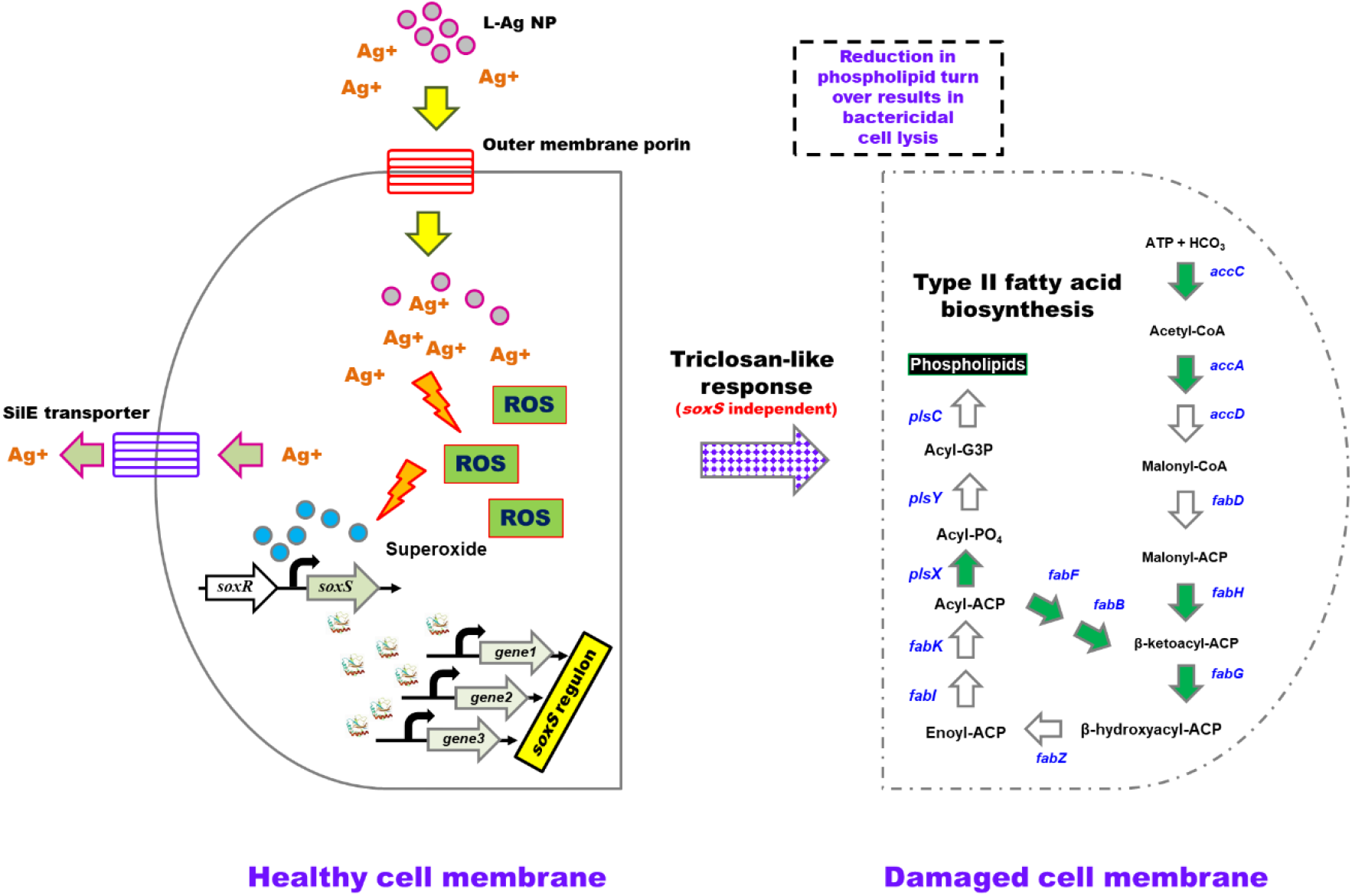
A model depicting the bactericidal action of L-Ag NPs on *K. pneumoniae* MGH78578. L-Ag NPs and silver ions (Ag^+^) enter the bacterial cell *via* outer membrane porins. Increasing cytoplasmic Ag^+^ concentration elicits the production of reactive oxygen species (ROS) including superoxide radicals, which then activate the *soxS*-mediated oxidative stress response. The SilE efflux system functions to eliminate Ag^+^ from the cytoplasm. RNA-seq data identified 7 genes (denoted by the green arrowheads) associated with type II fatty acid biosynthesis that are down-regulated leading to reduced phospholipid biosynthesis. This reduction in phospholipid turnover will affect the membrane stability leading to cell lysis underpinning the bactericidal activity. Grey arrows represent genes that are activated. White arrows represent genes that are either not identified in the *K. pneumoniae* MGH78578 genome or not differentially regulated when the bacteria are exposed to L-Ag NPs.

### The type II fatty acid biosynthesis genes were down-regulated in L-Ag NP exposed *K. pneumoniae* MGH78578

Since *K. pneumoniae* MGH78578 expressed an MDR phenotype,(13) L-Ag NP exposure dependent activation of *acrAB-tolC, marRAB,* and *sil* genes should, in principle, confer silver resistance in *K. pneumoniae*. Additionally, the strong oxidative stress response should enable the bacterium to counter the oxidative stress induced by L-Ag NPs. Both of these mechanisms should render *K. pneumoniae* MGH78578 resistant against silver. However, *K. pneumoniae* MGH78578 was found to be susceptible to L-Ag NPs. We hypothesized that the bactericidal activity of L-Ag NPs could be due to a hitherto unknown mechanism, rather than the most commonly reported oxidative stress based bactericidal activity.(27) To investigate, we compared our RNA-seq data to the transcriptional response of *E. coli* exposed to 37 antimicrobial compounds.(63) This global transcriptional profile of *E. coli* exposed to 37 antibiotics generated a list of 447 genes whose signature expression pattern characterized the mechanism of action associated with each antibiotic, leading the authors to hypothesize that the antibacterial action mechanism of any unknown/uncharacterized compounds can be deduced by comparing the transcriptional profile of a compound of interest with that of these 447 *E. coli* MG1665 K12 genes. We compared our exposed *K. pneumoniae* MGH78578 transcriptional data set with each individual dataset obtained from *E. coli* global transcriptional profile. Our comparative analysis showed that the highest number of similarly expressed gene pairs were obtained from the <u>triclosan exposure</u> dataset, suggesting that exposure to L-Ag NPs induced a *triclosan-like* exposure response in *K. pneumoniae* MGH78578. Some 76% of the gene pairs (73/96) at 5 min L-Ag NP exposure (Dataset WS3 and WS4) and 74% (87/117) at 30 min exposure (Dataset WS5 and WS6) had similar expression patterns compared with the *E. coli* triclosan dataset.

Triclosan is a broad-spectrum antimicrobial compound that acts by inhibiting FabI (a NADH-dependent enoylacyl carrier), a protein belonging to the type II fatty acid biosynthesis, part of a well-conserved pathway that is essential for bacterial survival. Inhibition of fatty acid biosynthesis compromises the bacterial cell membrane. During exposure to L-Ag NPs, some *K. pneumoniae* MGH78578 genes associated with type II fatty acid biosynthesis including *fabA/H/D/G/F/B* were significantly downregulated particularly during the adaptive response at 30 min. The downregulation of the different *fab* genes signals a compromised type II fatty acid biosynthesis underpinning the *triclosan-like* L-Ag NP bactericidal activity. Importantly, *fadL* and *fadD* were downregulated. FadL is a porin that transports extracellular fatty acids across the outer membrane to the inner membrane where they are activated by acyl CoA synthetase FadD. Downregulation of both *fadL* and *fadD* shows that extracellular fatty acids are not transported leading to subsequent suppression of type II fatty acid biosynthesis pathway. We, however, did not observe any differential expression in the *fabI* gene, possibly because the triclosan affects FabI post-translationally.

Recently, the TraDIS-Xpress approach involving a transposon mutant library and massively parallel sequencing of transposon chromosome junctions was used to identify *E. coli* genes that respond to triclosan exposure.(64) In comparison with these data, we identified genes that were both enriched in the *E. coli* TraDIS-Xpress dataset and that were found to be differentially expressed in our *K. pneumoniae* MGH78578 dataset. Common genes like *purL, purH, waeL, wzx* were downregulated at either one or both time points while *metB* was upregulated. Similarly, *K. pneumoniae* MGH78578 genes like *trkA, pcnB, infB,* and *ubiB/F* were downregulated at least in one or both time points, while *lonH* was upregulated (Dataset WS1). TraDIS-Xpress selected *rbs* in their triclosan exposure screen. In *E. coli,* RbsABC forms the ABC-type high-affinity D-ribose transporter, while RbsD/K phosphorylates D-ribose to D-ribose 5-phosphate.(65) Though we did not observe a statistically significant differential expression for *rbsB,* down-regulation was observed for *rbsD* and *rbsC,* showing similarly compromised D-ribose uptake. These common observations across different datasets gave confidence to our earlier observation that L-Ag NP exposure elicits a *triclosan-like* bactericidal effect in *K. pneumoniae* inhibiting type II fatty acid biosynthesis.

### The L-Ag NP based antibacterial action is cumulative of oxidative stress and fatty acid biosynthesis inhibition

We investigated whether the L-Ag NP based antibacterial effect was primarily due to type II fatty acid biosynthesis inhibition or as a cumulative effect of both oxidative stress and fatty acid biosynthesis inhibition. We exposed the isogenic *K. pneumoniae MGH78578 ΔsoxS* mutant, which has a compromised oxidative stress response mechanism,(54) to L-Ag NPs. The MIC of L-Ag NPs reduced to 15 μg mL^−1^ from 21 μg mL^−1^ showing that oxidative stress did contribute to a significant antibacterial effect. The antibacterial effect of L-Ag NPs is, therefore, a cumulative effect of both type II fatty acid biosynthesis inhibition and oxidative stress responses.

### Concluding observations

L-Ag NPs were very efficient in killing MDR *K. pneumoniae* MGH78578, triggering oxidative stress and a *triclosan-like* mechanism to exert their anti-*Klebsiella* effect. With toxicity studies in appropriate cell models, L-Ag NPs may be a future candidate as a novel antimicrobial agent. We hypothesized that our RNA-seq data could provide clues as to how *K. pneumoniae* might develop resistance against L-Ag NPs. Genes including *rbsD/C, trkA, pcnB* and *infB* were downregulated during the adaptive response at 30 min. Transposon insertional mutants in *E. coli* genes *trkA, pcnB* and *infB* exhibit better survival in the presence of triclosan.(64) Further, transposon insertion in *rbsB,* inhibiting the periplasmic ribose-binding domain of the RbsABC ribose importer, gave a selective survival advantage in the presence of triclosan. We hypothesize that by downregulating these genes during the adaptive response, *K. pneumoniae* MGH78578 could elicit strategies to combat the effects of exposure whilst resistance against L-Ag NPs. Exposure to sub-inhibitory concentrations of antimicrobial compounds is one of the main drivers in the evolution of AMR. By continuous exposure to sub-inhibitory concentrations of silver and the subsequent downregulation of these genes, *K. pneumoniae* might offer a transient resistance to L-Ag NPs. This hysteresis effect might be the prelude to developing a fully expressed antimicrobial resistance mechanism against silver.

## Materials and Methods

### Bacteria and media

*Klebsiella pneumoniae* MGH78578 (ATCC^®^700721) is a clinical isolate chosen for its MDR phenotype(47, 66) and the availability of its whole genome sequence (NC_009648.1). Bacteria were grown in Luria Bertani (LB) and modified LB (mLB) media constituted without NaCl.(39, 67) Bacteria were recovered from long-term storage (at −80°C in glycerol stocks) in mLB medium (casein enzyme hydrolysate 10 g L^−1^ and yeast extract 5 g L^−1^, pH 7.2 ± 0.2) for 12 h at 37 °C with shaking (150 rpm). For the preparation of inoculum, an overnight grown bacterial culture was inoculated into freshly prepared mLB medium and grown until the mid-log phase (OD_600 nm_ 0.5 to 0.6). An overnight grown bacterial culture (approximately 10^7^ CFU mL 1) was inoculated separately in freshly prepared LB and mLB broth medium. Bacterial growth was determined by measuring the optical density (OD) at 600 nm at 2 h time intervals. Biological experiments were carried out in both technical and biological duplicates.

### Synthesis of L-Ag NPs

L-Ag NPs were synthesized following a protocol adapted from Ashraf et al (see SI).(31) To determine the size, shape, crystallinity and surface capping, L-Ag NPs were characterized by Transmission Electron Microscopy (TEM), Energy Dispersive Spectroscopy (EDS), X-ray Diffraction (XRD), UV-vis spectroscopy, Fourier Transform Infrared spectroscopy (FTIR). The zeta potential of L-Ag NPs was measured with a Zetasizer Nano ZS (Malvern Instruments). The acellular oxidative potential of L-Ag NPs was assessed by measuring the depletion in antioxidants (uric acid, ascorbic acid, reduced glutathione) by HPLC following incubation of L-Ag NPs in a simplified synthetic respiratory tract lining fluid for 4 h at 37 °C.(68) The dissolution kinetics of L-Ag NPs in mLB was measured by inductively coupled plasma optical emission spectrometry (ICP-OES) (Avio 200, PerkinElmer). All measurements were performed in duplicate. The detailed protocols are described in the supplementary information (SI) file. Freshly prepared L-Ag NPs were used for all the experiments.

### Antibacterial susceptibility determinations

The minimum inhibitory concentration (MIC) was determined using the broth microdilution assay following CLSI guidelines. Bacterial cells were exposed to L-Ag NPs (ranging from 0.5 to 64 μg (Ag) mL^−1^) and incubated at 37 °C in the dark for 24 h. Bacterial growth was determined by measuring the OD_600 nm_. In addition, the free lysozyme was tested in parallel for any antibacterial activity. Bacterial cells exposed to L-Ag NPs were spread plated and incubated for 12 h at 37 °C to determine the minimum bactericidal concentration (MBC). Media without L-Ag NPs and bacterial cells were used as positive- and negative-controls respectively.(67) All experiments were done in biological duplicates and the results are represented as mean ± standard deviation.

### Mode of action studies-measurement of reactive oxygen species (ROS)

The generation of ROS inside the bacterial cell following exposure to L-Ag NPs was measured using a 2,7-dichlorodihydrofluorescein diacetate (DCFH-DA) assay.(69) DCFH-DA was added to the bacterial cell suspension at a final concentration of 10 μM and incubated for 1 h at 37 °C in the dark. Free dye was then separated from the DCFH-DA loaded bacterial cells by centrifugation at 8,000 rpm for 15 min followed by washing with PBS. Bacterial cells were exposed to different (sub)-MIC concentrations of L-Ag NPs (i.e. MIC_25_, MIC_50_, MIC_75_, and MIC_100_) for 30 min at 37 °C in fresh mLB medium with shaking (150 rpm). The fluorescence intensity of dichlorodihydrofluorescein (DCF) was detected using a Perkin Elmer VICTOR X Multilabel Plate Reader (USA) at an excitation and emission wavelength of 485 and 535 nm respectively. The experiments were performed in duplicates and the results expressed as mean ± standard deviation.

The effect of L-Ag NPs on the bacterial cell envelope was examined by TEM. To prepare sample for TEM analysis, bacterial cells exposed to MIC_75_ L-Ag NPs were centrifuged at 8,000 rpm for 10 min. Subsequently, the supernatant was discarded, and the bacterial pellet was washed twice with PBS followed by fixation in 2.5% v/v electron microscopy grade glutaraldehyde in 0.05 M sodium cacodylate buffer pH 7.2 for 2 h at 4 °C. TEM samples were prepared by drop-casting the bacterial cell suspension on to a carbon-coated copper grid that was later imaged on a Hitachi H-7650 TEM instrument (Hitachi High-Technologies Corporation) at an acceleration voltage of 100 kV. Bacterial cells that were not exposed to L-Ag NPs were used as the control. A minimum number of 50 bacterial cells were analyzed on different TEM images in each condition.

The membrane damage was analyzed using the MDA and anthrone assays. Bacterial cells were treated with different (sub)-MIC concentrations of L-Ag NPs, i.e. MIC_25_, MIC_50_, MIC_75_, and MIC_100_. The determination of MDA concentration was done as described previously.(70) The anthrone assay was done as described previously.(67) The experiments were performed in biological duplicates and the results expressed as mean ± standard deviation.

The intracellular concentration of Ag was measured by ICP-OES following exposure to MIC_75_ L-Ag NPs for 5, 30 and 60 min at 37 °C. A freshly grown bacterial culture (approximately 10^7^ CFU mL^−1^) was exposed to MIC_75_ L-Ag NPs followed by incubation for 5, 30 and 60 min at 37 °C in mLB. After incubation, bacterial cells were pelleted by centrifugation at 8,000 rpm for 10 min at 4 °C then dried and digested in a 1 mL mixture of H_2_O_2_:HNO_3_ (50:50) for 2 h. After acid digestion, the final volume was made up to 10 mL and filtered using 0.22 μm syringe filter. The silver concentration of the sample was measured by ICP-OES (Avio 200, PerkinElmer, USA).(38) Untreated bacterial cells were taken as a control for the respective time points. The experiments were performed in biological duplicates and the results expressed as mean ± standard deviation. Detailed protocols are described in SI.

### RNA sequencing

A freshly grown culture of *K. pneumoniae* MGH78578 from mid-log10 phase (approximately 10^7^ CFU mL^−1^) was exposed to MIC_75_ L-Ag NPs for 5 and 30 min at 37 °C in mLB broth. Bacterial cells not exposed to L-Ag NPs were selected as a control for each time point. RNA was isolated using RNAeasy extraction kit (Qiagen) and treated with Turbo DNase kit (Ambion’s). RNA integrity was assessed using a Bioanalyzer 2100 RNA 6000 nanochip (Agilent Technologies). RNA library sequencing was performed at the Centre for Genomic Research, University of Liverpool, UK. Ribosomal RNA was removed with a Ribo-Zero rRNA removal kit (Illumina). Libraries were prepared with NEBNext Directional RNA Library Prep Kit (BioLabs). Pooled libraries (Table S1) were loaded on cBot (Illumina) and cluster generations was performed. Single-sequencing using 150 bp read length was performed on lane of the HiSeq 4000 sequencer (Illumina). Raw sequencing data was processed as described in SI. The RNAseq data produced from the present work were deposited to the NCBI-GEO database and are available under the accession number GSE151953.

### Comparative gene expression analysis with E. coli MGH1655 exposed to individual antibiotics

A comparison of our transcriptional data with the global transcriptome profiling of *E. coli* MGH1665 K12 strain exposed to 37 antibiotics was carried out. Only those genes that gave statistically significant differential expression patterns were selected from both datasets. Our gene expression dataset was compared to each of the 37 *E. coli* transcriptional profiles and genes that had similar expression patterns (both up-and down-regulation) were counted.

#### Validation of RNA-seq data by quantitative RT-PCR analysis

RNA isolated from the samples were converted to cDNA by using high-capacity RNA to cDNA kit (Thermo fisher, Ireland). Primers based on the selected genes of interest were designed with 6-FAM/ZEN/IBFQ double-quenched probes and synthesized commercially by Integrated DNA Technologies (IDT, Belgium) (Table S2). The cDNA was used as a template and analysis was done by the addition of PrimeTime Gene Expression Master Mix. qRT-PCR was performed in an Eppendorf Mastercycler realplex ep gradient S (Eppendorf, UK). This analysis was carried out in two biological replicates each along with three technical replicates. The fold-change in the expression of the genes of interest was determined by the method of Livek and Schmittgen(71), i.e. 2^−ΔΔCt^ method using *rho* as a housekeeping gene.

#### Statistical analysis

The results for the biochemical assay were analysed using unpaired Student t-test as appropriate for the dataset. The qRT-PCR measurements data were statistically analysed using Prism software (v. 8.0 GraphPad Software) following the two-way analysis of variance. Boneferroni method was used to analyse the multiple comparisons. The symbol ‘ns’ used in the graphs corresponds to statistically non-significant with p > 0.05. The asterisk symbols in the graphs correspond to * p ≤ 0.05, ** p ≤ 0.01, and *** p ≤ 0.001. All the data points represent the mean of two independent measurements. The uncertainties were represented as standard deviations. In RNAseq results, NSE represents non-significant expression.

## Acknowledgments

Authors are thankful to BITS Pilani, University College Dublin, and Université de Paris for providing lab facilities. VP acknowledges CSIR-SRF [09/719(0090)/2018-EMR-l] and EMBO Short Term Fellowship (ESTF 7654) for the financial support. This project was supported, in part, by the Ulysses PHC Scheme (CNRS). We acknowledge Justine Renault, Linh-Chi Bui, and the Bioprofiler Facility (BFA) for the HPLC analysis.

## Author Contributions

SD, SS, SF, JP designed the study. VP, AB, SD characterized the silver NPs. VP and SS carried out the RNA-seq experiments. SKS generated the bioinformatics based gene expression dataset. SS, SD, VP, SF carried out the detailed RNA-seq dataset analysis that led to the triclosan observation. All authors read and approved the manuscript.

