## Supporting Information for "Silver nanoparticles induce a *triclosan-like* antibacterial action mechanism in multi-drug resistant *Klebsiella pneumoniae*" (from another version of this preprint: version 1, 2020-10-25)

**Table of Content**

Legends for Datasets WS1 to WS6
Supplementary Methods
Figures S1 to S4
Tables S1 to S2
SI References

**Other supplementary materials for this manuscript:**

Datasets WS1 to WS6

**Legends for Datasets WS1 to WS6.**

**WS-1.** Differential expression of all *K. pneumoniae* MGH 78578 genes during exposure to L-Ag NPs at 5 and 30 min post-treatment.

**WS-2.** Comparison of L-Ag NP exposure RNAseq data with the 'Oxidative SoxS regulon' dataset (1).

**WS-3.** Comparison of *K. pneumoniae* MGH78578 L-Ag NP exposure gene expression dataset at 5 min post-treatment with *E. coli* exposed to 37 antimicrobials (2). Green and purple boxes denote up- and downregulated genes respectively. Only genes present in genomes of both organisms that generated statistically significant gene expression data in both datasets are considered for comparison. '1' means that the expression in *K. pneumoniae* MGH78578 agrees with the *E. coli* one when exposed to a particular antimicrobial compound. Otherwise denoted as '0'.

**WS-4.** Comparison of *K. pneumoniae* MGH78578 L-Ag NP exposure gene expression dataset with *E. coli* exposed to triclosan. The dataset was further compared with the *K. pneumoniae* MGH78578 'oxidative soxS regulon' dataset (1).

**WS-5.** Comparison of *K. pneumoniae* MGH78578 L-Ag NP exposure gene expression dataset at 30 min post-treatment with *E. coli* exposed to 37 antimicrobials (2). Green and purple boxes denote up- and downregulated genes respectively. Only genes present in genomes of both organisms that generated statistically significant gene expression data in both datasets are considered for comparison. '1' means the expression in *K. pneumoniae* MGH78578 agrees with the *E. coli* one when exposed to a particular antimicrobial compound. Otherwise denoted as '0'.

**WS-6.** Comparison of *K. pneumoniae* MGH78578 L-Ag NP exposure gene expression dataset at 30 min post-treatment with *E. coli* exposed to triclosan. The data was further compared with the *K. pneumoniae* MGH78578 'oxidative soxS regulon' dataset (1).

### Supplementary Methods

**Synthesis of lysozyme coated silver nanoparticles (L-Ag NPs).** Two mg lysozyme powder was added to 20 mL solution of 1 mM $AgNO_3$ with vigorous stirring at 120 °C whilst being continuously heated for 15-20 min. In this reaction mixture, lysozyme functions as both reducing and stabilizing agent. This step was then followed by the addition of 1 mL 1M NaOH and the heat reflux continued for 1 h at the same temperature. Synthesis of L-Ag NPs was visible by the development of light- to dark-red color of the solution (3, 4). Freshly prepared L-Ag NPs were used for all the experiments.

**Characterization of L-Ag NPs.**

UV-visible spectroscopy. The UV-vis spectrum of L-Ag NPs was measured in water on a V-630 spectrophotometer (Jasco Corporation, Japan). The appearance of the plasmon peak was characteristic of L-Ag NP synthesis.

Transmission Electron Microscopy (TEM) and Energy Dispersive X-Ray Spectroscopy (EDS). Samples for TEM analysis were prepared by drop-casting L-Ag NPs on carbon-coated copper grids followed by vacuum drying. High-resolution TEM images were recorded on a Hitachi H-7650 TEM instrument (Hitachi High-Technologies Corporation, Japan) at an acceleration voltage of 100 kV. Silver composition was confirmed by SAED pattern determination using a JEOL-2100 TEM instrument (JEOL, Japan). Elemental analysis of L-Ag NPs was carried out using a Quantax EDS attachment (Bruker, UK).

X-Ray Diffraction (XRD). The X-ray diffraction pattern of L-Ag NPs were observed on a Rigaku MiniFlex II Benchtop XRD System (Rigaku Company, USA) operated at a voltage of 45 kV and current of 40 mA with Cu Kα radiation. The crystal phase was analysed by comparing the calculated values of interplanar spacing and the corresponding intensities of diffraction peaks with the standard theoretical values of the Powder Diffraction File database (PCPDFWIN; JCPDS-ICDD 2008).

Zeta potential. The zeta potential of L-Ag NPs was measured in water and in mLB medium at 37 °C on a Zetasizer, Malvern Instruments, USA. The zeta-potential was calculated by fitting the electrophoretic mobility with the Smoluchowski model. The experiment was performed in triplicate and the results expressed as mean ± standard deviation.

Fourier Transform Infrared spectroscopy (FTIR). The FTIR spectra of L-Ag NPs were measured in water using a Shimadzu IR Prestige-21 FTIR spectrometer (Shimadzu, Japan). The measurements were carried out in the 400 – 4,000 $cm^{-1}$ range with a resolution of 4 $cm^{-1}$. The analysis was performed in duplicate.

Oxidative potential assay. The acellular oxidative potential of L-Ag NPs was assessed by measuring the depletion in antioxidants, uric acid, ascorbic acid, and reduced glutathione, following incubation of L-Ag NPs at a concentration of 1 to 10 μg $mL^{-1}$ in a simplified synthetic respiratory tract lining fluid (sRTLF) for 4 h at 37 °C. sRTLF composed of 200 μM uric acid (UA), 200 μM ascorbic acid (AA) and 200 μM reduced glutathione (GSH) in 9 g $L^{-1}$ NaCl at pH 7.4 was prepared following guidelines (5).Then the solutions were centrifuged on SpinX 0.2 μm cellulose acetate filters at 12,000 g to remove the NPs. The antioxidant (UA, AA, GSH) and oxidized glutathione (GSSG) concentrations were analyzed by reverse-phase HPLC on a Kromasil 100-5C18 column using 25 mM sodium phosphate and 0.5 mM octane sulfonic acid for the mobile phase A, and acetonitrile for the mobile phase B. The absorbance of UA, AA, GSH, and GSSG were measured at 280 nm, 243 nm, and 210 nm respectively. The depletion in reduced glutathione (GSH) was confirmed by the formation of oxidized glutathione (GSSG). A sample containing sRTLF only was used as a reference. The depletion in antioxidant was expressed as the ratio between the antioxidant concentration difference between the reference and the sample, divided by the antioxidant concentration of the reference. All the measurements were performed in triplicates.

**Preparation of bacterial culture.** Bacteria were recovered from long-term storage (at -80°C in glycerol stocks) in modified Luria Bertani (mLB) medium (casein enzyme hydrolysate 10 g $L^{-1}$ and yeast extract 5 g $L^{-1}$, pH 7.2 ± 0.2) for 12 h at 37 °C with shaking (150 rpm). For the preparation of inoculum, an overnight grown bacterial culture was inoculated into freshly prepared mLB medium and grown until the mid-log phase ($OD_{600\ nm}$ 0.5 to 0.6). Biological experiments were carried out in both technical and biological duplicates.

**Minimum inhibitory concentration (MIC) assay.** Bacterial cultures were grown to mid-log phase then diluted in fresh mLB to a final cell density of approximately $10^7$ CFU $mL^{-1}$. Initially, bacterial cells were exposed to a broad dilution range (consisting of two-fold dilutions) of 0.5 to 64 µg (Ag) $mL^{-1}$ L-Ag NPs and then incubated at 37 °C until early stationary phase (ESP) (6, 7). These dilution ranges were subsequently narrowed further, to obtain a precise MIC value. Bacterial growth was determined by measuring the $OD_{600\ nm}$. In addition, the free lysozyme was tested in parallel for any antibacterial activity. All experiments were done in biological duplicates and the results are represented as mean ± standard deviation.

**Minimum bactericidal concentration (MBC) assay.** Following exposure to L-Ag NPs, 100 µL of a bacterial sample was spread plated directly onto freshly prepared mLB agar plates. Inoculated plates were then incubated at 37 °C for 24 h and any growth recorded. Based on the results obtained using broad range concentrations, further experiments were performed to narrow the concentration range of L-Ag NPs and identify the precise MBC value. Media without L-Ag NPs and bacterial cells were used as positive- and negative-controls respectively (8).

**2,7-dichlorodihydrofluorescein diacetate (DCFH-DA) assay**. A fresh culture of bacterial cells (approximately $10^7$ CFU $mL^{-1}$) was obtained and washed three times with phosphate buffered saline (PBS) followed by the suspension in fresh mLB medium. DCFH-DA was added to the bacterial cell suspension at a final concentration of 10 µM and incubated for 1 h at 37 °C in the dark. Free dye was then separated from the DCFH-DA loaded bacterial cells by centrifugation at 8,000 rpm for 15 min followed by washing with PBS. Bacterial cells were exposed to different (sub)-MIC concentrations of L-Ag NPs (i.e. $MIC_{25}$, $MIC_{50}$, $MIC_{75}$, and $MIC_{100}$) for 30 min at 37 °C in fresh mLB medium with shaking (150 rpm). The fluorescence intensity of dichlorodihydrofluorescein (DCF) was detected using a Perkin Elmer VICTOR X Multilabel Plate Reader (USA) at an excitation and emission wavelength of 485 and 535 nm respectively. The experiments were performed in duplicates and the results expressed as mean ± standard deviation.

**Sample preparation for TEM analysis of bacterial cells.** To prepare sample for TEM analysis, bacterial cells exposed to $MIC_{75}$ L-Ag NPs were centrifuged at 8,000 rpm for 10 min. Subsequently, the supernatant was discarded, and the bacterial pellet was washed twice with PBS followed by fixation in 2.5% v/v electron microscopy grade glutaraldehyde in 0.05 M sodium cacodylate buffer pH 7.2 for 2 h at 4 °C. TEM samples were prepared by drop-casting the bacterial cell suspension on to a carbon-coated copper grid that was later imaged on a Hitachi H-7650 TEM instrument (Hitachi High-Technologies Corporation) at an acceleration voltage of 100 kV. Bacterial cells that were not exposed to L-Ag NPs were used as the control. A minimum number of 50 bacterial cells were analyzed on different TEM images in each condition.

**Malondialdehyde (MDA) assay**. Bacterial cells were treated with different (sub)-MIC concentrations of L-Ag NPs, i.e. $MIC_{25}$, $MIC_{50}$, $MIC_{75}$, and $MIC_{100}$. The determination of MDA concentration was done as described previously (9). The experiments were performed in biological duplicates and the results expressed as mean ± standard deviation.

**Anthrone assay**. To validate the damage to the bacterial cell membrane, an anthrone assay was carried out. In this assay, carbohydrates are hydrolysed into furfurals and hydroxyl-methyl furfurals, which further react with anthrone and form a blue-green complex that was detected at 620 nm using Jasco V-630 UV visible spectrophotometer (USA) (10). Estimation of carbohydrate released from the bacterial cytoplasm treated with the different (sub)-MIC concentration of L-Ag NPs, i.e. $MIC_{25}$, $MIC_{50}$, $MIC_{75}$, and $MIC_{100}$, was done as described previously (8). The experiments were performed in biological duplicates and the results expressed as mean ± standard deviation.

**Intracellular silver concentration in *K. pneumoniae* following treatment with L-Ag NPs.** A freshly grown bacterial culture (approximately $10^7$ CFU $mL^{-1}$) was exposed to $MIC_{75}$ L-Ag NPs followed by incubation for 5, 30 and 60 min at 37 °C in mLB. After incubation, bacterial cells were pelleted by centrifugation at 8,000 rpm for 10 min at 4 °C then dried and digested in a 1 mL mixture of $H_2O_2$:$HNO_3$ (50:50) for 2 h. After acid digestion, the final volume was made up to 10 mL and filtered using 0.22 µm syringe filter. The silver concentration of the sample was measured by ICP-OES (Avio 200, PerkinElmer, USA) (11). Untreated bacterial cells were taken as a control for the respective time points. The experiments were performed in biological duplicates and the results expressed as mean ± standard deviation.

**RNA sequencing data processing**. Raw RNAseq data were processed as described earlier (12) with few modifications. Briefly, RTA version 1.18.61 was used for the processing of raw sequencing data and CASAVA 1.8.4 was used to generate FASTQ-files. Sequence quality of the RNAseq reads were analysed using FastQC (https://www.bioinformatics.babraham.ac.uk/projects/fastqc/). Reads were aligned against reference genome of *K. pneumoniae* MGH78578 (NCBI RefSeq NC_009648.1) using Segemehl with default mapping parameters (13) and uniquely mapped reads were considered for the differential gene expression analysis. Read counts (number of reads that aligned to a specific gene) for each gene were quantified using custom Perl scripts. The counts were transformed using the VOOM function (14) in the Bioconductor package limma v3.26.1 (15) in R v3.5.2. Principle Component Analysis (PCA) was performed to analyse quality of the RNAseq libraries. Linear regression models within limma were used to identify differentially expressed genes between samples w/wo L-Ag NPs exposure. Finally, p-values for differential expression were adjusted with a false discovery rate (FDR, Benjamini-Hochberg) with a p-value $< 0.05$ as significant.

**Validation of RNA-seq data by quantitative RT-PCR analysis.** RNA isolated from the samples were converted to cDNA by using High-capacity RNA to cDNA kit (Thermo fisher, Ireland). Primers based on the selected genes of interest were designed with 6-FAM/ZEN/IBFQ double-quenched probes and synthesized commercially by Integrated DNA Technologies (IDT, Belgium) (Table S2). The cDNA was used as a template and analysis was done by the addition of PrimeTime Gene Expression Master Mix. qRT-PCR was performed in an Eppendorf Mastercycler realplex ep gradient S (Eppendorf, UK). This analysis was carried out in two biological replicates each along with three technical replicates. The fold-change in the expression of the genes of interest was determined by the method of Livek and Schmittgen (17), i.e. $2^{-\Delta\Delta Ct}$ method using *rho* as a housekeeping gene.

**Statistical analysis.** The results for the biochemical assay were analysed using unpaired Student t-test as appropriate for the dataset. The qRT-PCR measurements data were statistically analysed using Prism software (v. 8.0 GraphPad Software) following the two-way analysis of variance. Boneferroni method was used to analyse the multiple comparisons. The symbol 'ns' used in the graphs corresponds to statistically non-significant with $p > 0.05$. The asterisk symbols in the graphs correspond to * $p \leq 0.05$, ** $p \leq 0.01$, and *** $p \leq 0.001$. All the data points represent the mean of two independent measurements. The uncertainties were represented as standard deviations. In RNAseq results, NSE represents non-significant expression.

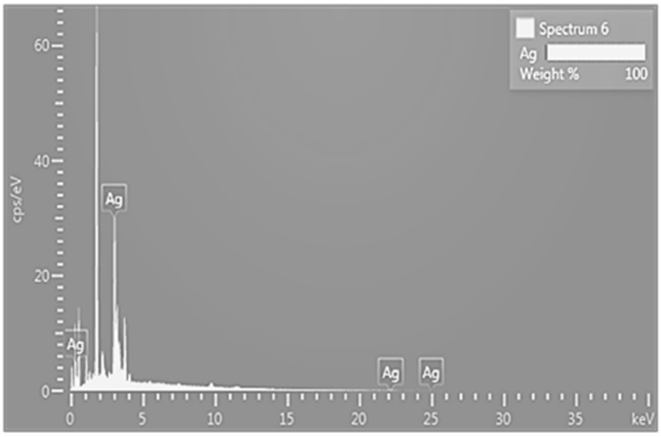


**Fig. S1.** TEM-EDS spectrum of L-Ag NPs deposited on a carbon copper grid.

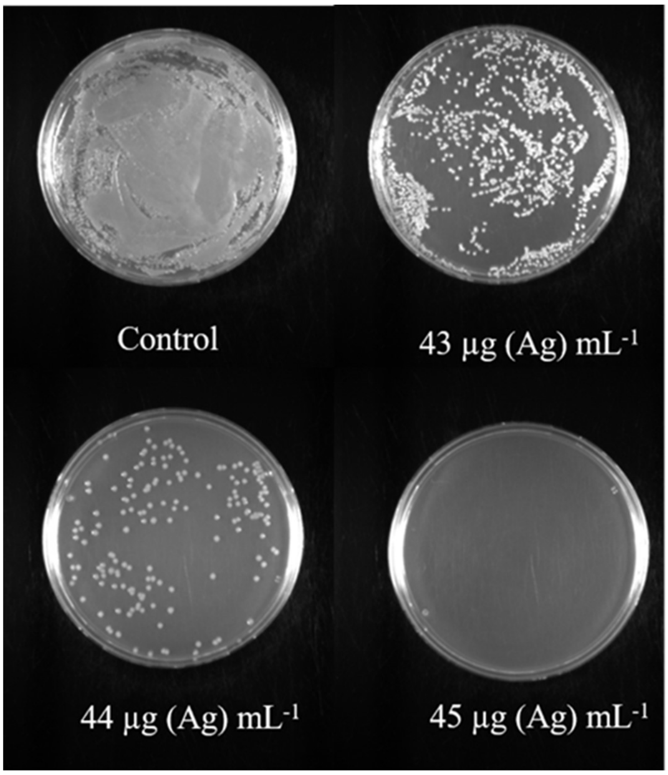


**Fig. S2.** Determination of the minimum bactericidal concentration (MBC) of L-Ag NPs against *K. pneumoniae* MGH78578 in mLB agar plates.

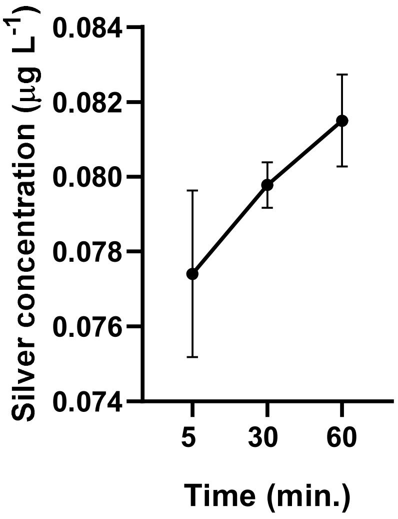


**Fig. S3.** Intracellular silver concentration measured in *K. pneumoniae* MGH78578 following exposure to $MIC_{75}$ L-Ag NPs for 5, 30, and 60 min at 37 °C in mLB.

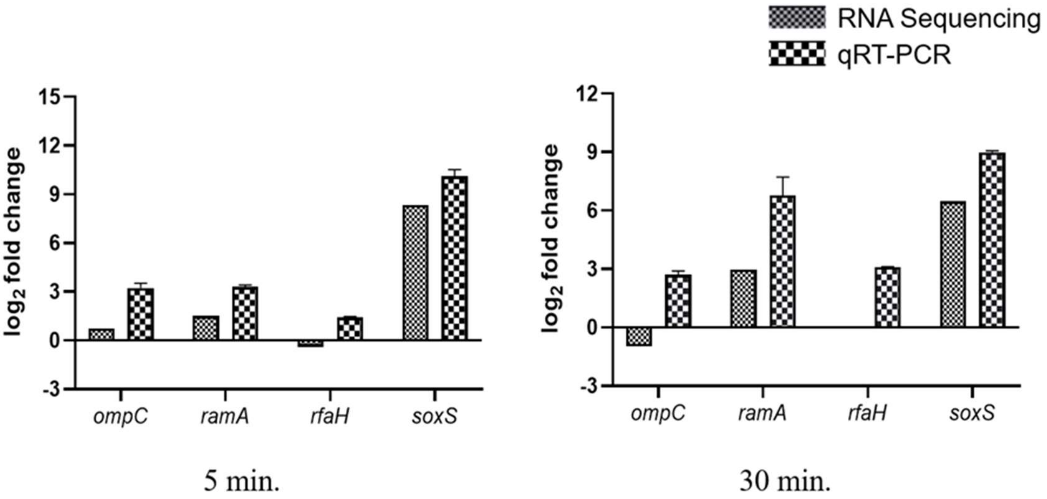


**Fig. S4.** Comparison of the RNA-seq and qRT-PCR data for *K. pneumoniae* MGH78578 exposed to $MIC_{75}$ L-Ag NPs at 5 and 30 min post-treatment.

**Table S1.** Overview of the RNA sequencing libraries for *K. pneumoniae* MGH78578.

| Sample no. | RNA-seq library and experimental condition | Treatment time |
|---|---|---|
| **1.** | Control 5 min. (Replicate 1) | **5 min.** |
| **2.** | Control 5 min. (Replicate 2) | |
| **3.** | $MIC_{75}$ L-Ag NP 5 min. (Replicate 1) | |
| **4.** | $MIC_{75}$ L-Ag 5 min. (Replicate 2) | |
| **5.** | Control 30 min. (Replicate 1) | **30 min.** |
| **6.** | Control 30 min. (Replicate 2) | |
| **7.** | $MIC_{75}$ L-Ag 30 min. (Replicate 1) | |
| **8.** | $MIC_{75}$ L-Ag 30 min. (Replicate 2) | |

**Table S2.** Details of gene specific primers used for qRT-PCR experiments on *K. pneumoniae* MGH78578.

| Sample no. | Target genes | Oligonucleotide primer DNA sequence | Amplicon size (bp) |
|---|---|---|---|
| **1.** | ***ompC*** | Forward: 5'-TGG TCG TTG ATC TGG GTT TC-3' | 99 |
| | | Reverse: 5'-AAT TGA CGG TCT GCA CTA CTT-3' | |
| | | **Probe**: /56-FAM/CAT GCG TGT/Zen/AGG CGT GAA AGG C/3IABkFQ/ | |
| **2.** | ***ramA*** | Forward: 5'-TTC CGC TCA GGT GAT TGA TAC-3' | 104 |
| | | Reverse: 5'-TTG CAG ATG CCA TTT CGA ATA C-3' | |
| | | **Probe**: /56-FAM/TAA CCT GCA/Zen/TCA ACC GCT GCG TAT/3IABkFQ/ | |
| **3.** | ***rfaH*** | Forward: 5'-TGC GTT TAC CGC GAA TGA-3' | 106 |
| | | Reverse: 5'-AAA CGT GGG CAG CTA CAA-3' | |
| | | **Probe**: /56-FAM/ACT GAC GTT/Zen/CCA GAT GTT CCT GGG/3IABkFQ/ | |
| **4.** | ***soxS*** | Forward: 5'-GCA TCA CGG TAC GGA ACA T-3' | 112 |
| | | Reverse: 5'-GAT GAA CAT ATC GAT CAA CCA CTT AAC-3' | |
| | | **Probe**: /56-FAM/TCG AGT ATC/Zen/CTG ACT TTC TGG CGA CT/3IABkFQ/ | |
| **5.** | ***rho*** | Forward: 5'-CTC AGG AAG AGC TGC AGA AA-3' | 108 |
| | | Reverse: 5'-CTT GGT CAT TGC CAG CTT ATT G-3' | |
| | | **Probe**: /56-FAM/CGA TGG GTG/Zen/AGA TTG ATG CGA TGG A/3IABkFQ/ | |
